# A cell-based screen identifies HDAC inhibitors as activators of RIG-I signaling

**DOI:** 10.1101/2021.09.28.462211

**Authors:** Eugenia Fraile-Bethencourt, Marie H Foss, Dylan Nelson, Sanjay V Malhotra, Sudarshan Anand

## Abstract

Enhancing the immune microenvironment in cancer by targeting the nucleic acid sensors is becoming a potent therapeutic strategy. Among the nucleic acid sensors, activation of the RNA sensor Retinoic Acid-inducible Gene (RIG-I) using small hairpin RNAs has been shown to elicit powerful innate and adaptive immune responses. Given the challenges inherent in pharmacokinetics and delivery of RNA based agonists, we set out to discover small molecule agonists of RIG-I using a cell-based assay. To this end, we established and validated a robust high throughput screening assay based on a commercially available HEK293 reporter cell line with a luciferase reporter downstream of tandem interferon stimulated gene 54 (ISG54) promoter elements. We first confirmed that the luminescence in this cell line is dependent on RIG-I and the interferon receptor using a hairpin RNA RIG-I agonist. We established a 96-well and a 384-well format HTS based on this cell line and performed a proof-of-concept screen using an FDA approved drug library of 1200 compounds. Surprisingly, we found two HDAC inhibitors Entinostat, Mocetinostat and the PLK1 inhibitor Volasertib significantly enhanced ISG-luciferase activity. This luminescence was substantially diminished in the null reporter cell line indicating the increase in signaling was dependent on RIG-I expression. Combination treatment of tumor cell lines with Entinostat increased RIG-I induced cell death in a mammary carcinoma cell line that is resistant to either Entinostat or RIG-I agonist alone. Taken together, our data indicates an unexpected role for HDAC1,-3 inhibitors in enhancing RIG-I signaling and highlight potential opportunities for therapeutic combinations.

## Introduction

Mammalian cells have mechanisms that activate the innate immune response to provide immediate defense upon infection. Infections are recognized by pattern recognition receptors (PRRs), which identify pathogen-associated molecular patterns (PAMPs) ^1^. PRRs that identify pathogen DNA or RNA in the cytoplasm are known as cytosolic nucleic sensors. One of the most important PRRs for viral infections is the retinoic acid-inducible gene I (RIG-I). RIG-I recognizes short double stranded RNA (dsRNA) with di or tri-phosphate group at the 5’end ^2^. RIG-I, encoded by DDX58 (DEADbox helicase 58 gene) in humans, has two caspase activation and recruitment domains (CARDs) in the N-terminus, a central helicase domain (HD) and C-terminal domain (CTD) ^3^. Under normal conditions, CARDs are bound to the HD in an auto-repress conformation. If viral RNA is detected, RIG-I dimerizes, and CARDs are released to bind with the CARDs from the mitochondrial activator of virus signaling (MAVS) protein. RIG-I and MAVS interaction triggers activation of the interferon regulatory factor (IRF) 3 and NF-κB. Activated IRF3 and NF-κB translocate to the nucleus to activate the expression of type 1 interferon (IFN-I) and subsequently, proinflammatory and anti-viral genes. IFN-I acts in an auto and paracrine manner activating a number of ISGs, whose products regulate cell growth, metabolism and immune response ^4^.

Several studies have identified synthetic RIG-I agonists which trigger IFN-I response for use as vaccine adjuvants, to treat viral infection or to enhance the immunogenicity in cold tumors ^5,6^. In fact, RIG-I agonists are showing promising results in pre-clinical and clinical trials. For instance, RNA loop SLR14 improved the effects of anti-PD1 in an MC-38 tumor model ^7^; and MK-4621 is showing promising results in clinical trials in solid tumors (NCT03065023 and NCT03739138). However, delivery challenges as well as pharmacokinetics of synthetic RNA agonists argue for the development of small molecule agonists to target RIG-I. Thus, we decided to develop a cell-based high throughput screening (HTS) assay. Here we have demonstrated that it is feasible to identify small molecule agonists of RIG-I using this assay and discovered unexpected crosstalk between HDAC, PLK1 signaling and the RIG-I pathway.

## Results

### An ISG-luciferase reporter cell line is a sensitive tool to study RIG-I activation *in vitro*

HEK-Lucia™ RIG-I (Invivogen) cells have been engineered with the DDX58 (RIG-I human gene) and a secreted Lucia luciferase gene under the ISG54 promoter. We first established that the luciferase signal in the cell lysates were equivalent to the secreted luciferase signal in the supernatants and therefore decided to assay the cell lysates to minimize liquid handling (data not shown). We found that RIG-I activation by recognition of 3p-hpRNA (RIG-I agonist) triggers IFN-I response, which activates luciferase expression (Sup. Fig. 1A). In contrast HEK-Lucia™ Null cells which bear the same ISG54 responsive luciferase gene but lack DDX58 had significantly decreased luminescence upon RIG-I agonist treatment but retained equivalent responsiveness to IFN-β treatment (Sup. Fig. 1B). Importantly, addition of an IFNAR inhibitor decreased the luminescence induced by the RIG-I agonist suggesting the reporter activity is dependent on IFN signaling (Sup. Fig. 1C). To optimize the RIG-I activity assay, we performed dose-response experiments in 96 well plates following the manufacturer’s indications (approximately 5×10^5^ cell/well) at 24h and 48h. We observed a robust dose dependent increase in luciferase signal with the RIG-I agonist (Fig. 1). Z’ values were 0.68 for 24 hours (Fig. 1A) and 0.88 for 48 hours (Fig. 1B) at 0.3 μg/ml of RIG-I or control agonist. We optimized cell number and kinetics and found that at 6h, luciferase expression was detected from 5,000 cells/ well (Fig 1C); while at 48 hours, only 500 cells/well were needed to detect luminescence (Fig. 1B). Our data indicate that this 3p-hpRNA is a potent RIG-I activator, and the HEK-Lucia™ RIG-I as a sensitive tool to test RIG-I activation *in vitro*.

**Figure 1.**
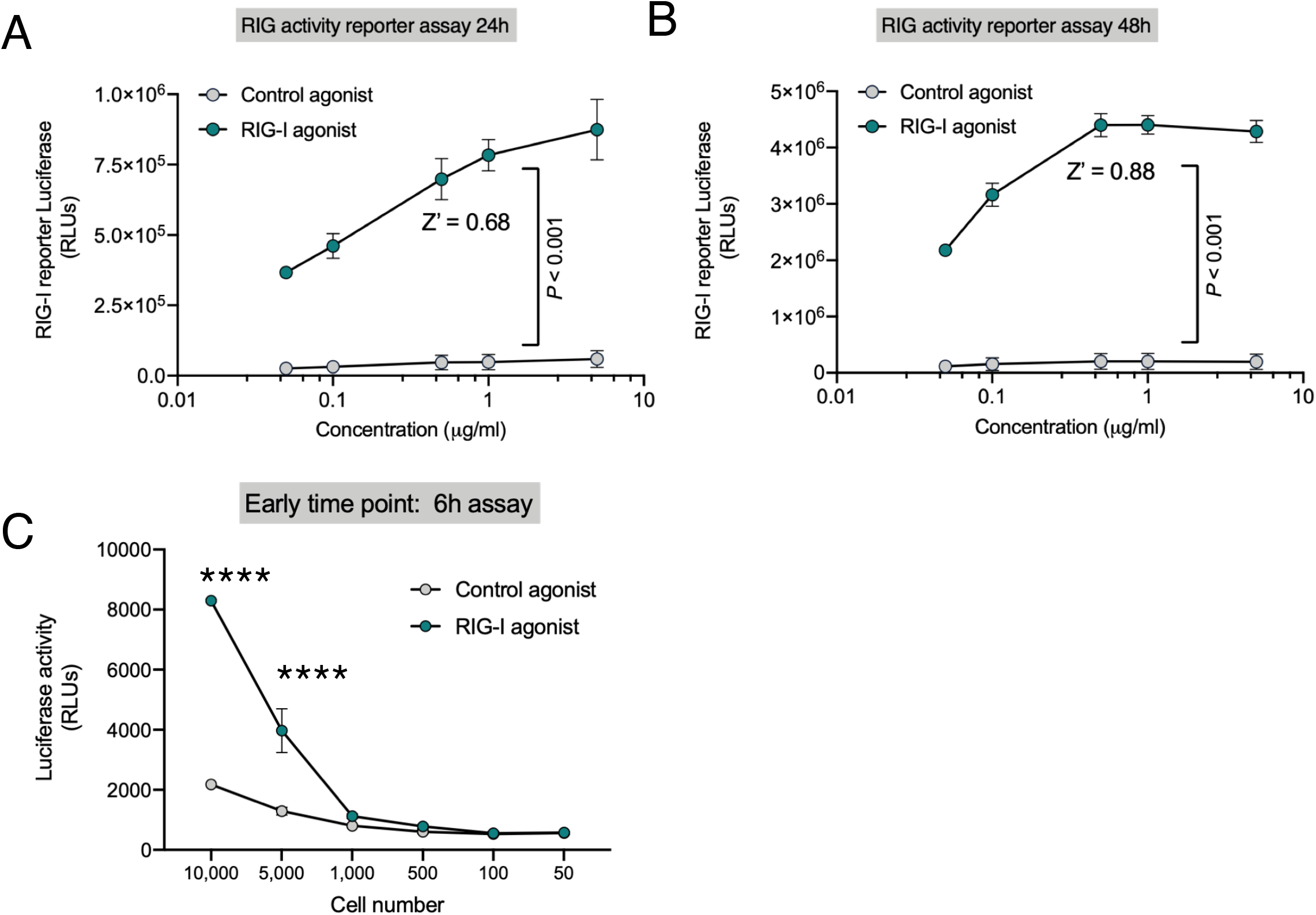
Sensitive Reporter Assay for measuring RIG-I activity. HEK293 RIG-I cells were seeded onto 96-well plates and treated with the indicated concentrations of a control agonist or RIG-I agonist. Luciferase activity was measured in supernatants as per manufacturer’s recommendations at A) 24h B) 48h and C) 6h with different cell numbers. P value from a two-way ANOVA with post-hoc Sidak’s correction or unpaired two-tailed Student’s T-test.

### High Throughput Screening identifies RIG-I activity in an FDA approved drug library

To find new compounds that could activate RIG-I, we carried out a HTS using an FDA approved drug library. This library included 1,430 compounds in water or DMSO, with well characterized bioactivity, bioavailability, and safety profiles. We confirmed that the assay yielded robust signal in 384-well plates and tolerated DMSO with no decrease in luminescence till 5% DMSO (Figure 2A). We tested the FDA library compounds in 8-point 1/3 Log_10_ dose curves from 10 mM to 3 μM over several days of independent runs. Results indicated that our assay worked robustly with an average Z’ of 0.57 (Figure 2B), good signal to background noise difference (S/B) above 65 (Figure 2C). We defined a ‘hit’ as any compound with luminescence of at least 5 SDs above the average background of DMSO treated wells. We found that 12 compounds in the FDA library achieved this threshold in our HTS (Figure 2D).

**Figure 2.**
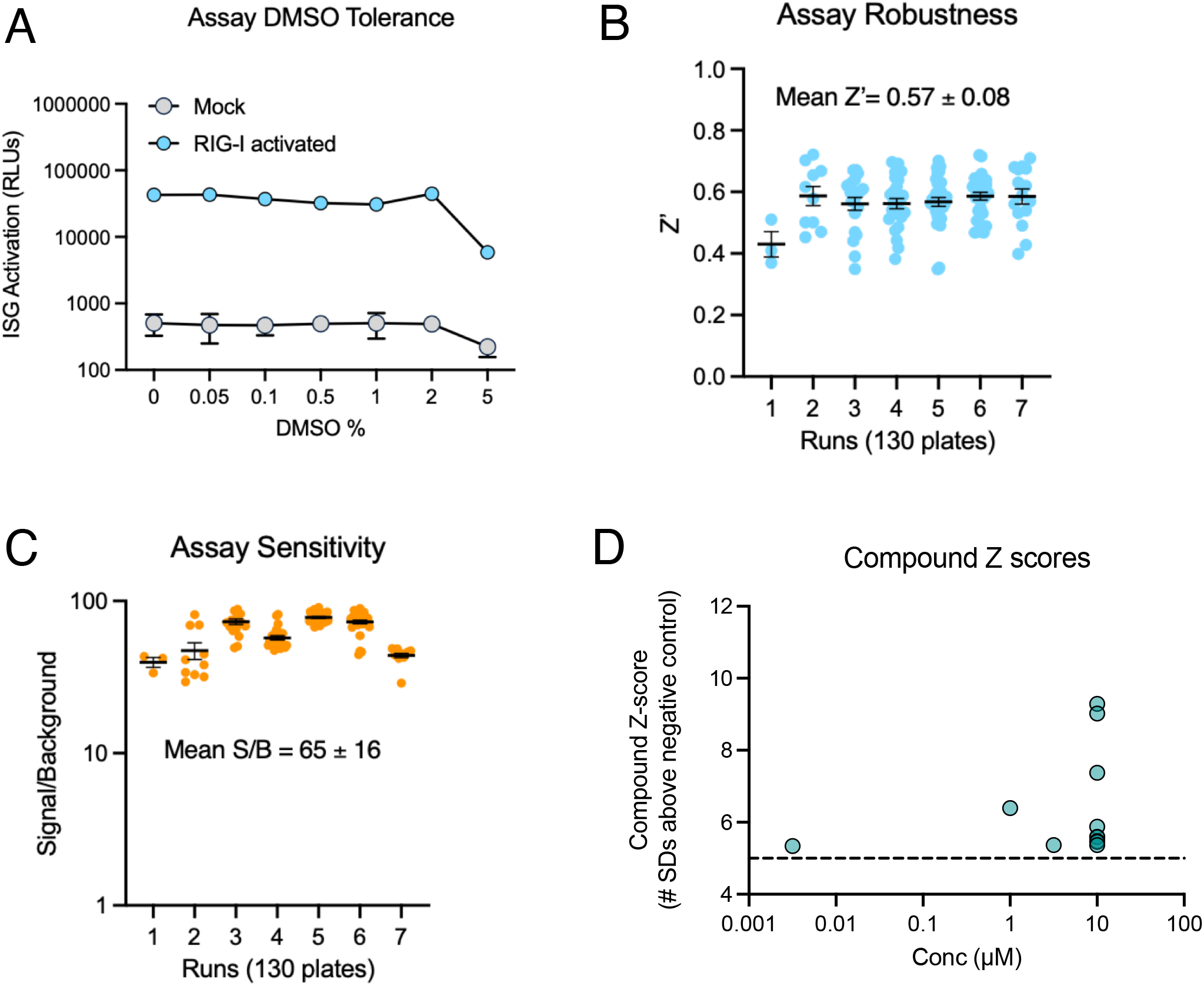
Miniaturization and validation of RIG activity HTS assay in 384 well plate. A) DMSO tolerance of HEK293 RIG-I cell line as measured by luciferase activation with positive control RIG-I agonist B-C) Luciferase activity was measured in cell lysates as per manufacturer’s recommendations at 24h post treatment with three compound libraries across 8 doses-the FDA approved drug collection, a LOPAC collection and a collection of natural product derived compounds from Microsource. Screening data from 7 independent runs totaling 130 plates show B) high Z’ values, C) good signal to noise ratios. D) Compound Z-scores of hits (at any concentration) in the FDA library defined as number of standard deviations (SDs) above the mean of negative control wells.

Interestingly, we found three compounds-2 HDAC inhibitors-Entinostat, Mocetinostat and the PLK1 inhibitor Volasertib increased the ISG activation signal at more than one dose (Figure 3A-C). IFN-I response can be activated by different pathways in the cells. In fact, HEK-Lucia™ RIG-I and HEK-Lucia™ Null cells endogenously express NOD1, TLR3 and TLR5. However, our data showed that HEK-Lucia™ RIG-I cells do not respond to LPS or Poly I:C across a range of concentrations (Sup. Fig. 2). To study the RIG-I specificity of these compounds, we assayed them in HEK-Lucia™ Null cells for responses across 4-log doses. We found that for the HDAC inhibitors and PLKi, and to a lesser extent the JAK inhibitor Fediratinib, the luminescence was dependent on expression of RIG-I in the cells (Figure 3D). Independent dose response experiments confirmed that the activity of Entinostat, Mocetinostat and Volasertib was specific to the RIG-I expressing but not null cells (Figure 3E-G).

**Figure 3.**
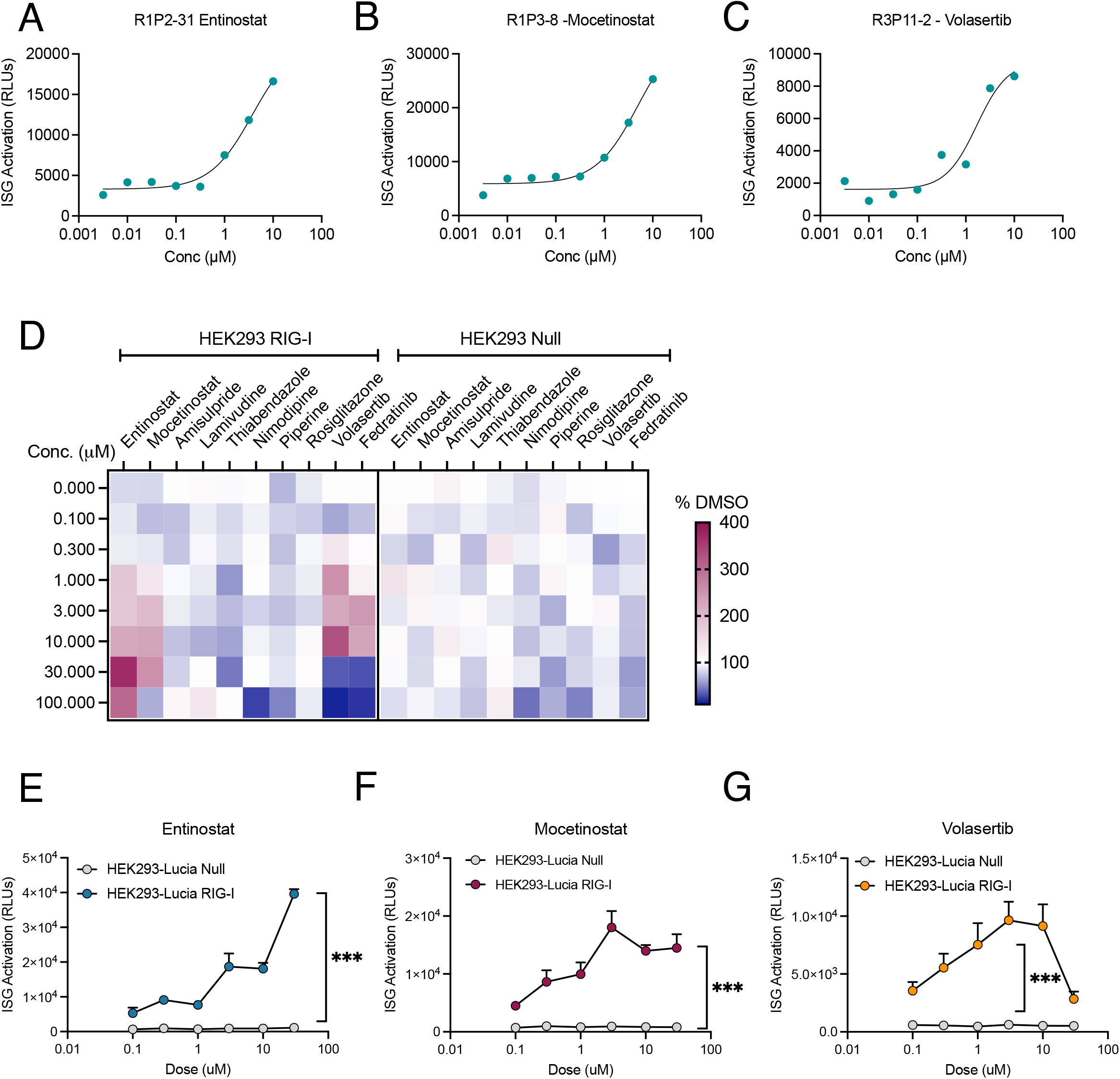
HDAC inhibitors and a PLK inhibitor increase ISG activation in a RIG-I dependent manner. Dose response curves showing low μM activation of ISG-luciferase by A-B) HDAC inhibitors Entinostat and Mocetinostat and C) PLK1 inhibitor volasertib. D) Heatmap depicts % change vs DMSO treatment for top 10 hits in the same cell line. E-G) Comparison of dose dependent ISG-Luciferase activation by the drugs in Null vs RIG-I expressing reporter cell lines. *** P<0.001 by 2-way ANOVA with Sidak’s post-hoc correction.

### Entinostat enhances RIG-I signaling in a resistant tumor cell line *in vitro*

Since we established that these compounds activate RIG-I signaling in HEK293 cells, we next evaluated their ability to drive cell death in tumor cell lines. Both HDAC inhibitors as well as the PLK inhibitor are approved drugs with potent anti-tumor activity in a range of cancers. To evaluate whether these inhibitors can confer additive or even synergistic effects with RIG-I agonists, we chose the 4T1 mammary carcinoma cells which showed resistance to both RIG-I agonist induced cell death (Figure 4A) and Entinostat induced cell death (Figure 4B). We found that combination of RIG-I agonist with Entinostat increased apoptosis compared more significantly to either RIG-I alone or the drug alone (Figure 4C). While the mechanisms by which Entinostat increases RIG-I cell death remains to be determined, we observed two bona fide type I interferon response genes, Mx1 and Usp18 were significantly induced by Entinostat treatment.

**Figure 4.**
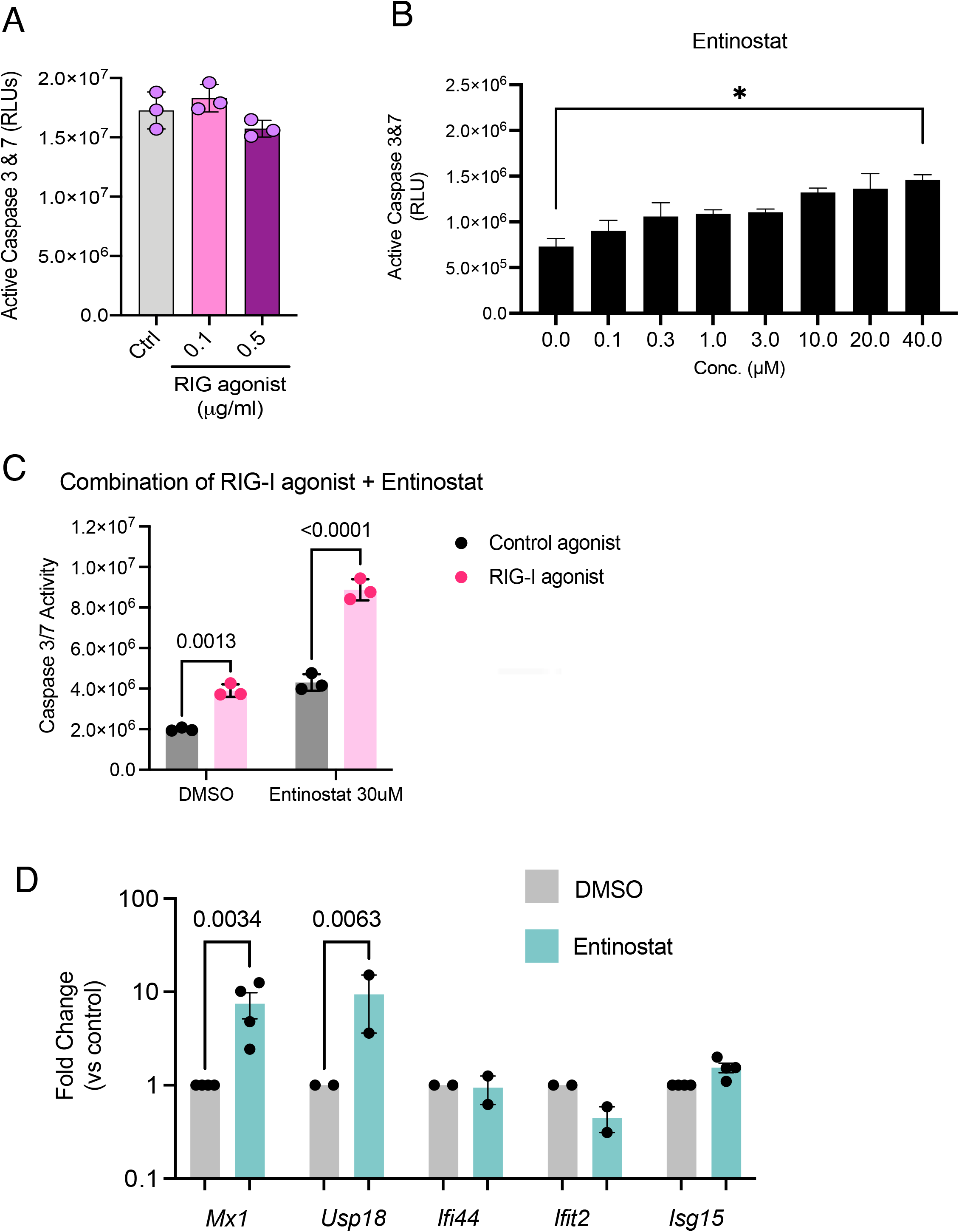
Activation of RIG-I signaling in tumor cell lines in response to HDAC and PLK inhibitors. A) 4T1 mammary carcinoma cells were treated with RIG-I agonist alone or (B) Entinostat alone or (C) RIG-I agonist at 0.5 ug/ml followed by Entinostat at the indicated concentration. 24h later cell death was measured by a CaspaseGlo assay. 4T1 mammary carcinoma cells were treated with DMSO, Entinostat (30uM). qRT-PCR for indicated genes is shown. P values from a two-way ANOVA with a post-hoc Sidak’s test (A-C) or an uncorrected Fisher’s LSD test (D).

Taken together, our data indicates that a cell-based screening assay can function robustly to perform high throughput screening for RIG-I agonists. We surprisingly observed that RIG-I signaling can be activated by HDAC inhibitors and a PLK1 inhibitor and showcase a potential utility in driving cell death in tumors that maybe resistant to RIG-I or HDAC inhibition monotherapies.

## Discussion

RIG-I is one of the most important defenses against viral infection in cells functioning as a PRR, but emerging data suggests broader roles in cancer. Previous studies showed RIG-I as tumor suppressor in different cancer types such as hepatocellular carcinoma and glioblastoma multiforme ^8,9^. Several studies have shown that RIG-I activation is a potent driver of immune responses in cancer ^7,10–12^. While RNA agonists have shown remarkable activity in preclinical models and a phase I clinical trial, there are still challenges in delivering RNA to solid tumors in humans over longer durations of treatment ^13^. Therefore, we set out to establish a screening assay for the discovery of small molecules that can activate RIG-I in a selective manner. Our data indicates that this cell-based assay is robust and highly sensitive to identify compounds that activate RIG-I dependent ISG signaling. We also serendipitously discovered that HDAC inhibition and PLK inhibition can also activate RIG-I signaling. Observations from independent experiments using our RIG-I null cell lines, the IFNAR inhibitors and the TLR ligand treatment experiments suggest that the luminescence in our assay is a) directly dependent on the ectopic expression of RIG-I b) is type I interferon dependent and c) is not driven by other typical RNA sensors in cells. We also show that the ability of HDAC inhibitors and PLK inhibitor to drive RIG-I signaling holds true in tumor cell lines in vitro. Interestingly these three inhibitors confer an additive benefit in driving cell death in combination with RIG-I activation in a cell line that is typically insensitive to RIG-I induced cell death.

While further in-depth studies are needed to validate the cross-talk between the RIG-I and HDAC/PLK pathways in tumor cells, we propose a few hypotheses on how these pathways interact. There is evidence that HDAC3 inhibition can increase transcription of type I interferons ^14^. Interestingly, it is thought that HDAC inhibition can also enhance the transcription of endogenous retroviral elements and other non-coding RNAs both in the human and mouse genome ^15^. Given the role of RIG-I in sensing RNA, it is possible that these endogenous RNAs can activate RIG-I. RIG-I is also regulated by acetylation at multiple sites. There is evidence that HDAC6 interacts with and acetylates RIG-I ^16,17^. Similarly, it has been reported that PLK1 can directly bind and phosphorylate MAVS leading to an increase in type I interferon signaling ^18^. Based on our data, we suspect distinct mechanisms are contributing to how these pathways can enhance RIG-I signaling.

Beyond the biological basis for how these important pathways engage and interact, there is also a potential clinical utility for our findings. Although HDAC inhibitors and PLK1 inhibitors have been extensively studied, their proposed mechanism(s) of action have been tumor centric ^19,20^. However, recent evidence suggests that HDAC inhibition may have roles in the tumor microenvironment ^21^. For instance, it has been shown that HDAC inhibition potentiates immunotherapy responses in CT26 and MC38 tumor models ^22^. Our observations introduce the possibility that HDAC inhibition also contribute type I interferons to activate the immune microenvironment. In summary, we have described a new, highly sensitive cell-based assay to discover small molecule agonists of RIG-I signaling and highlight a putative role for HDAC inhibition in activating RIG-I pathway in tumor cells. We anticipate further work will clarify both the biological mechanisms and the functional significance of this interaction in greater detail.

## Materials and Methods

### Cell culture

RIG-I reporter cell line HEK-Lucia™ RIG-I (Invivogen, #hkl-hrigi) was cultured in Dulbecco’s Modified Eagles Medium (DMEM), High Glucose (HyClone™) supplemented with 10% Fetal Bovine Serum (FBS) (Bio-Techne R&D Systems #S11550H), 30 μg/ml blasticidin, 100 μg/ml Normocin™ and 100 μg/ml of Zeocin™. HEK-Lucia™ Null cells (Invivogen, #hkl-null) were used as control and cultured in the same media with neither blasticidin nor Zeocin™. MC-38 (Kerafast #ENH204-FP) and CT-26 (ATCC #CRL-2638) 4T1 cells were cultured in DMEM or RPMI 1640 with L-Glutamine (Lonza™ BioWhittaker™ # BW12115F12) supplemented with 10% of FBS. Cell lines were routinely tested and found to be mycoplasma free. Cell line identities were genetically validated through Cell Line Authentication Service at OHSU Genomics Core Facility.

### RIG-I activity assay

HEK-Lucia™ RIG-I or Null cells were transferred into antibiotic free media (DMEM supplemented with 10% FBS) and seeded in 96 or 384 well plate at 70-85% confluence. RIG-I agonist (Invivogen #tlrl-hprna) or control agonist (Invivogen #tlrl-3prnac) were transfected with LyoVec™ (Invivogen #lyec-12) following the manufacturer’s instructions. Cells were incubated for 6-48 hours at 37ºC and 5% of CO^2^. QUANTI-Luc™ Gold was dissolved in sterile water and added directly into the wells. Luminescence was immediately measure in Spark® Multimode Microplate Reader (Tecan) set at 0.1 second of reading time.

### RNA extraction, RT-PCR and gene expression

Genomic RNA from cells and homogenized tumors was extracted using the GeneMATRIX Universal RNA Purification Kit (EuRx #E3598) following the manufacturer’s instruction. Samples were treated with DNAse I (Invitrogen™ #18068015) to avoid DNA contamination. Reverse transcription was performed with 500ng of RNA using the High-Capacity cDNA Reverse Transcription Kit (Cat: 4368814, Applied Biosystems). Gene expression was assayed by quantitative PCR using TaqManTM Master Mix II no UNG (Cat 4440049, Thermofisher Scientific) with the following probes: Ddx58 (Mm01216853_m1), Mx1 (Mm00487796_m1), Isg15 (Mm01705338_s1), Usp18 (Mm01188805_m1), Ifit2 (Mm00492606_m1), Ifi44 (Mm00505670_m1), Gapdh (Mm99999915_g1). Data was normalized to internal control Gapdh and gene expression quantified using the using the 2^−ΔΔCt^ method with the control treatment as reference.

### Cell Titer-Glo/Caspase Glo

Cells were seeded in white 96 well plates (Greiner Bio-One™ CellStar™) and treated for 24 hours. Cell Titer-Glo (Promega, #G7572) and Caspase 3/7 Glo (Promega #G8091) were added and assayed according to the manufacturer’s instructions. Luminescence was measure using either a Promega GloMax, BioTex Synergy 4 or Tecan Spark reader.

### High-throughput screening

Reporter cells were seeded in cell culture 384 well plates (Greiner, #781080) at 1.5×10^4^ cells/ well in 50 μl of DMEM supplemented with 10% FBS using BioTek™ MultiFlo™ FX Microplate Dispensers (BioTek Instruments). Cells were treated with an FDA approved library (Selleckchem) using a Sciclone ALH3000 Liquid Handler (PerkinElmer). Each compound was assayed across 8 doses and incubated for 24 hours. Using the Matrix WellMate dispenser (Thermo Scientific), 30 μl of QUANTI-Luc™ Gold was added. Immediately, luminescence was measured at 0.1 second of reading time. Results were analyzed using Dotmatics software.

For each plate, the Z’ factor was calculated as 1- (3 x SD positive control + 3 x SD of negative control)/|Mean of positive control -Mean of negative control). The Signal to Background ratios were calculated for each plate based on the mean signal from positive control wells over the mean signal of all the negative control wells.

### Statistics

All statistical analysis was performed using Prism software (GraphPad Software, San Diego, CA). Differences between pairs of groups were analyzed by Student’s *t*-test. Comparison among multiple groups was performed by one-way ANOVA followed by a post hoc test (Tukey’s or Holm-Sidak). In the absence of multiple comparisons, Fisher’s LSD test was used. Values of *n* refer to the number of experiments used to obtain each value. For experiments where the data was not normally distributed, we used the Kruskal-Wallis test. Values of *p* ≤ 0.05 were considered significant.

## Data and Material Availability

All data are contained within the manuscript. This study did not generate any unique materials.

## Acknowledgements

This work was supported by funding from NHLBI to S.A. (R01 HL137779 and R01 HL143803). We thank Sokchea Khou, Rebecca Ruhl, Adrian Baris for technical help and members of the Anand lab and Malhotra lab for useful discussions.

## Author Contributions

E. F-B, M.F., D.N, S.M. and S.A designed experiments, E. F-B and M.F. performed experiments, E. F-B, M.F., D.N, S.M. and S.A analyzed the data, E. F-B and S.A wrote the manuscript. All authors reviewed and edited the manuscript.

---

**Supplementary Figure 1.**
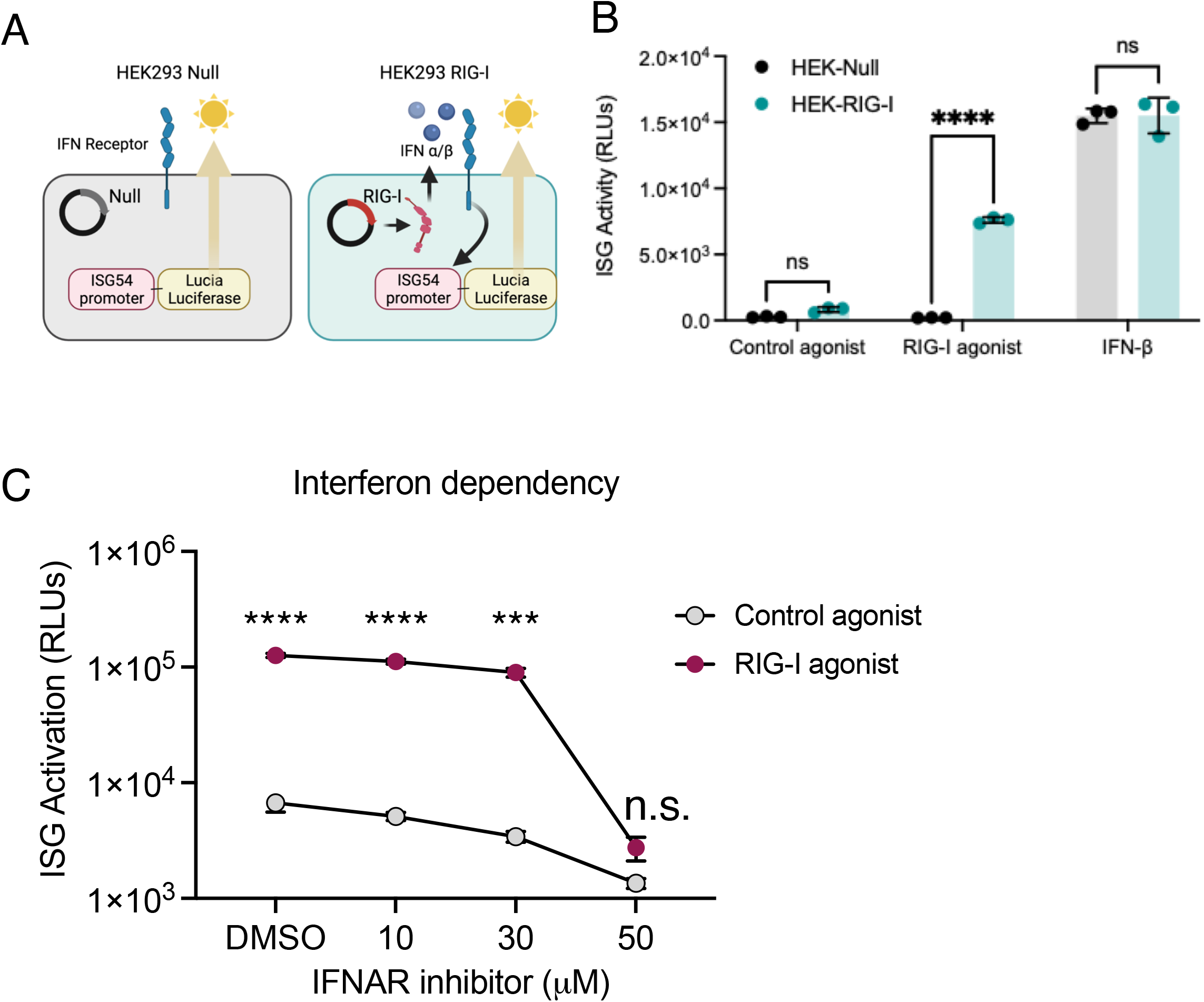
RIG-I and IFN dependent activation of an ISG54 luciferase reporter cell line. A-B) HEK293 reporter cell lines with an ISG54 responsive luciferase gene and a null vector or RIG-I expressing vector were treated with a control agonist or RIG-I agonist or IFN-β as a positive control. ISG activity was assessed using luminescence in cell lysates at 24h in the null cell line and the RIG-I expressing cell lines. ****P<0.0001 using ANOVA with post-hoc Sidak’s test. C) HEK293 RIG-I cells were treated with a control agonist or RIG-I agonist in the presence of indicated concentrations of an interferon alpha receptor antagonist. ISG activity was assessed by luminescence at 24h. ****P<0.0001 using Welch’s T-test with post-hoc Bonferroni-Sidak’s test.

**Supplementary Figure 2.**
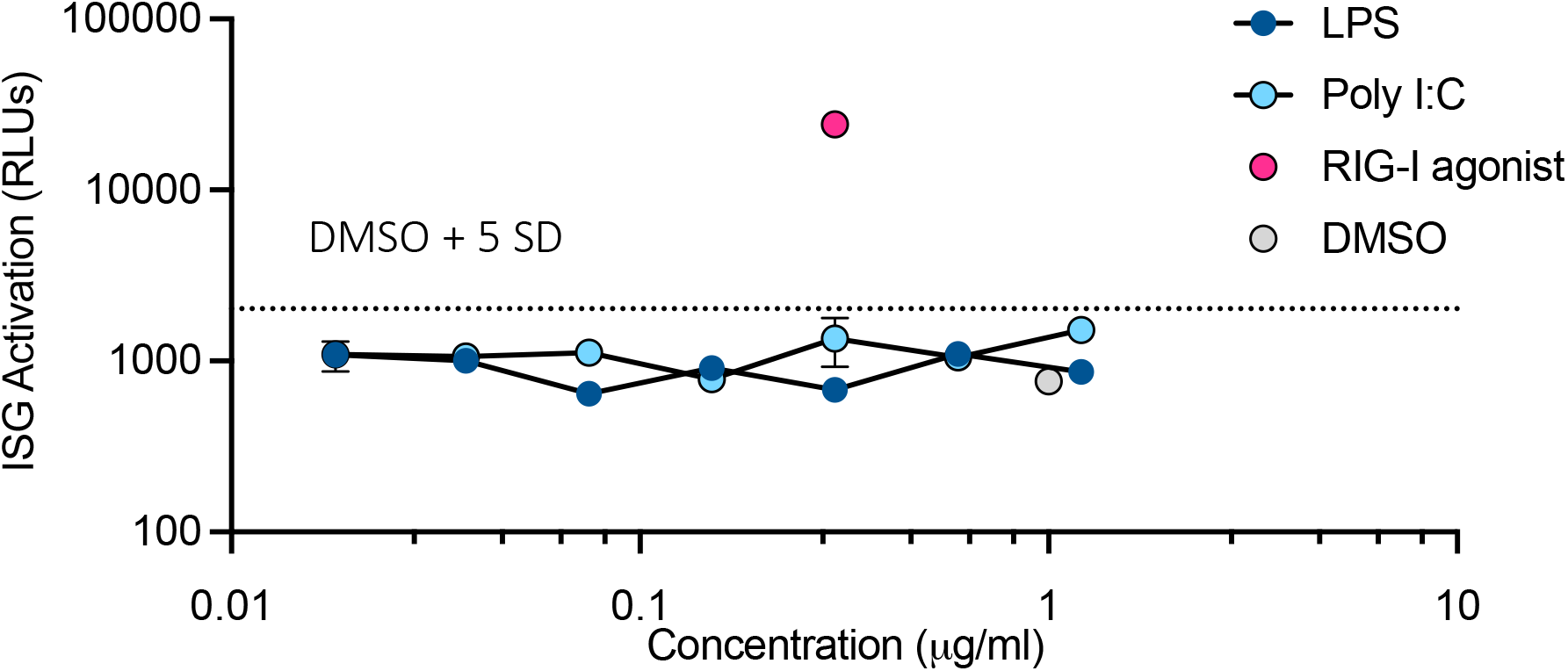
TLR ligands do not activate ISG54 luciferase in our reporter assay. HEK293 reporter cell lines were treated with different Poly I:C, TLR 3 agonist or LPS a TLR4 and TLR2 agonist at the indicated concentrations. RIG-I agonist was used at 0.3 ug/ml as a positive control and 1% DMSO as negative control. Dotted line indicates our hit criteria at 5 SD above the mean of negative control wells.

